# Cyclic and Multilevel Causation in Evolutionary Processes

**DOI:** 10.1101/830422

**Authors:** Jonathan Warrell, Mark Gerstein

## Abstract

Many models of evolution are implicitly causal processes. Features such as causal feedback between evolutionary variables and evolutionary processes acting at multiple levels, though, mean that conventional causal models miss important phenomena. We develop here a general theoretical framework for analyzing evolutionary processes drawing on recent approaches to causal modeling developed in the machine-learning literature, which have extended Pearl’s ‘do’-calculus to incorporate cyclic causal interactions and multilevel causation. We also develop information-theoretic notions necessary to analyze causal information dynamics in our framework, introducing a causal generalization of the Partial Information Decomposition framework. We show how our causal framework helps to clarify conceptual issues in the contexts of complex trait analysis and cancer genetics, including assigning variation in an observed trait to genetic, epigenetic and environmental sources in the presence of epigenetic and environmental feedback processes, and variation in fitness to mutation processes in cancer using a multilevel causal model respectively, as well as relating causally-induced to observed variation in these variables via information theoretic bounds. In the process, we introduce a general class of multilevel causal evolutionary processes which connect evolutionary processes at multiple levels via coarse-graining relationships. Further, we show how a range of ‘fitness models’ can be formulated in our framework, as well as a causal analog of Price’s equation (generalizing the probabilistic ‘Rice equation’), clarifying the relationships between realized/probabilistic fitness and direct/indirect selection. Finally, we consider the potential relevance of our framework to foundational issues in biology and evolution, including supervenience, multilevel selection and individuality. Particularly, we argue that our class of multilevel causal evolutionary processes, in conjunction with a minimum description length principle, provides a conceptual framework in which identification of multiple levels of selection may be reduced to a model selection problem.

## 1 Introduction

Causality is typically invoked in accounts of evolutionary processes. For instance, for a variant to be subject to direct selection, it is necessary that it has a causal impact on fitness. The role of causality is made explicit in axiomatic accounts of evolution [32]. Further, the formal framework of Pearl’s ‘do’-calculus [30] has been used explicitly in analyzing Mendelian Randomization [24], the relationship between kin and multilevel selection [28], and information derived from genetic and epigenetic sources in gene expression [13]. A number of features of evolutionary processes, however, limit the potential for direct formalization in the ‘do’-calculus framework, which requires causal relationships to be specified by a directed acyclic graph (DAG), and cannot represent causal processes at multiple levels. In contrast, cyclical causal interactions are ubiquitous in natural processes, for instance in regulatory and signaling networks which lead to high levels of epistasis in the genotype-phenotype map [38]. Further examples of cyclical causal interactions arise through environmental feedback, both in the generation of traits, leading to an extended genotypeenvironment-phenotype map [16], and across generations in the form of niche construction [21]. Hierarchy is also ubiquitous in evolution, and many phenomena, such as multicellularity and eusociality, seem to require a multilevel selection framework for analysis, implicitly invoking causal processes at multiple levels [27]. Such a framework would also seem necessary in analyzing major transitions in evolution [6].

A number of frameworks have been proposed in the machine-learning literature for extending Pearl’s ‘do’-calculus to allow for cyclic causal interactions. These include stochastic models with discrete variables [17], and deterministic [25] and stochastic [33] models with continuous variables. Further, approaches have been introduced for analyzing causal processes at multiple levels using the ‘do’-calculus [7, 33]. In [33], both of these phenomena are related through the notion of a *transformation*, which is a mapping between causal models which preserves causal structure. Coarse-graining is a particular kind of transformation, special cases of which involve mapping a causal model over micro-level variables into one over macro-level variables, and mapping a directed causal model which is extended across time into a cyclical model which summarizes its possible equilibrium states (subject to interventions).

In addition, the ‘do’-calculus has been combined with information theory in order to define notions of information specifically relevant to causal models, such as *information flow* [2], *effective information* [15], *causal specificity* [13] and *causal strength* [19]. Although not explicitly cast in causal terms, there has also been much interest in defining non-negative multivariate decompositions of the mutual information between a dependent variable and a set of independent variables, which may be collectively described as types of *Partial Information Decomposition* (PID) [3, 13, 40]. Such definitions, however, can only be applied in causal models with a DAG structure, leaving open the question of how causal information should be defined and decomposed in a system with cyclic interactions.

Motivated by the above, we propose a general causal framework for formulating models of evolutionary processes which allows for cyclic interactions between evolutionary variables and multiple causal levels, drawing on the transformation framework of [33] as described above. Further, we propose a causal generalization of the Partial Information Decomposition (Causal Information Decomposition, or CID) appropriate for such cyclic causal models, and show that our definition has a number of desirable properties and can be related to previous measures of causal information. We analyze a number of specific evolutionary models within our framework, including first, a model with epigenetics and environmental feedback, and second, a model of multilevel selection which we apply to the particular cases of group selection and selection between mutational processes in cancer. We analyze the CID in the context of both models, and demonstrate the conditions under which bounds can be derived between components of the CID and components of the PID associated with the observed distribution. Finally, we discuss the causal interpretation of Price’s equation and related results in our causal framework. In general, our analysis is intended both to help clarify conceptual issues regarding the role of causation in the models analyzed, as well as to aid in the interpretation of data when the assumptions of these (or similar) models are adopted, via the bounds introduced.

We begin in Sec. 2 by providing an overview of the issues related to cyclic and multilevel causality in evolution which motivate our approach, and informally introduce the Discrete Causal Network (DCN) and Causal Information Decomposition (CID) frameworks which form the basis of subsequent analyses (full definitions are given in Appendix A). Sec. 3 then outlines our general framework for Causal Evolutionary Processes (CEP). Sec. 4 and Sec. 5 analyze specific models of epigenetics with environmental feedback and multilevel selection within this framework respectively, and Sec. 6 provides a causal interpretation of Price’s equation and related results. Sec. 7 then concludes with a discussion, including the potential relevance of our framework to foundational issues in the philosophy of biology and evolution.

## 2 Cyclic and Multilevel Causality in Biology

We begin by outlining and motivating some of the basic concepts that will be used to develop our framework. We do so here in an informal way: technical definitions and proofs are given in Appendix A.

### Cyclic Causality

The *do*-calculus provides a compelling formalization of the mathematical structure of causation [30]. However, a requirement of this framework is that causal relationships between variables must form a Directed Acyclic Graph (DAG). A DAG is any graph (a collection of nodes and edges) which contains no directed loops (cycles). For instance, if gene A regulates gene B, and gene B regulates genes C and D (for example, the genes are transcription factors, and regulatory relationships are established via promoter binding), variables corresponding to the expression levels of these genes can be arranged in a graph with no cycles. Performing manipulations on gene B (*do*-operations) will thus affect the expression of genes C and D but not A. Pearl’s calculus provides exact rules for deducing the distribution of all variables after an intervention, which will always return a valid distribution; formally, the graph is altered by cutting all incoming edges to the manipulated node, and the joint distribution is recalculated using a delta distribution to represent the intervention.

However, such well-ordered networks are the exception rather than the rule in gene regulatory networks (GRNs). For instance, we could add to the above network a feedback interaction by assuming that gene D regulates A. Here, although A causes B’s expression locally, B also causes A’s expression via D. In fact, in this case no harm is done assuming the system has a solution as a whole, since any intervention will either split the cycle (genes A, B and D), or have no effect on it (gene C), and hence all distributions are well defined. Alternatively though, we could consider starting with the original graph, and in addition let C regulate D and D regulate C. Now, by intervening on B we are not guaranteed to find a solution for every intervention, even if the original system has a solution. Recent work has investigated the extension of Pearl’s calculus to graphs with cycles (where the graph manipulations are identical to the acyclic case), characterizing the situations in which solutions exist, and allowing systems to be defined which have a restricted set of interventions allowed [17, 25, 33]. A particular example which has a clear solution is a deterministic network governed by linear differential equations: 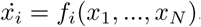, where the *x*_*i*_’s may be gene expression values for instance. Here, ‘solutions’ are taken to be the equilibrium points of the system, and these exist for any intervention provided the original system of equations *f*_1_, …*f*_*N*_ is *contractive*, meaning that it maps a given region of state-space to one with a smaller volume with time [33]. Stochasticity can be added as long as the functions are contractive almost surely.

In these examples, we have considered feedback processes in gene regulatory networks. However, similar feedback processes can occur at many levels. For instance, consider a psychological trait, such as depression. This trait may be ultimately caused by numerous aspects of brain structure and gene expression patterns; however, since we have pharmacological interventions which can control the severity of depression, these can introduce a feedback process from the environment to the molecular layer. In general, the expression of any trait may be subject to environmental feedback in this way.

We now note some features about the above. First, we have phrased both the GRN and the environmental feedback example in terms of a process in time. One way to deal with such cyclic structures is to ‘unroll’ them over time, that is, consider a discretized set of time points, and repeat all variables at each time, while connecting a given variable, say a gene, to its regulatory parents at the previous time-step. This will automatically generate an acyclic causal graph. The cyclic causal system corresponding to this temporal process can be considered to be that formed by the equilibrium distributions (assuming they exist) after certain ‘macro’ interventions are applied, which fix a particular variable to a given value across all times. The cyclic system can thus be considered a *coarse-graining* of the temporal acyclic system. Not all cyclical causal systems can be formed this way (see [33]), but our emphasis will be on such systems, given their prevalence in biology (although the framework is agnostic to the system’s origins). For convenience, in setting up our framework, we use a ‘Discrete Causal Network’ model (DCN, Appendix A; see also Fig. 1A for the relationships between all models in the paper), which allows cycles and variables which take discrete values, and is parameterized by a set of *probability kernels*, one for each variable, specifying the conditional distribution of the variable on the values of its graphical parents. We further introduce dynamic-DCN and equilibrium-DCN models (Appendix A, Def. 2.6), corresponding respectively to an underlying temporal process and coarse-grained equilibrium cyclic causal model as discussed above.

**Fig. 1.**
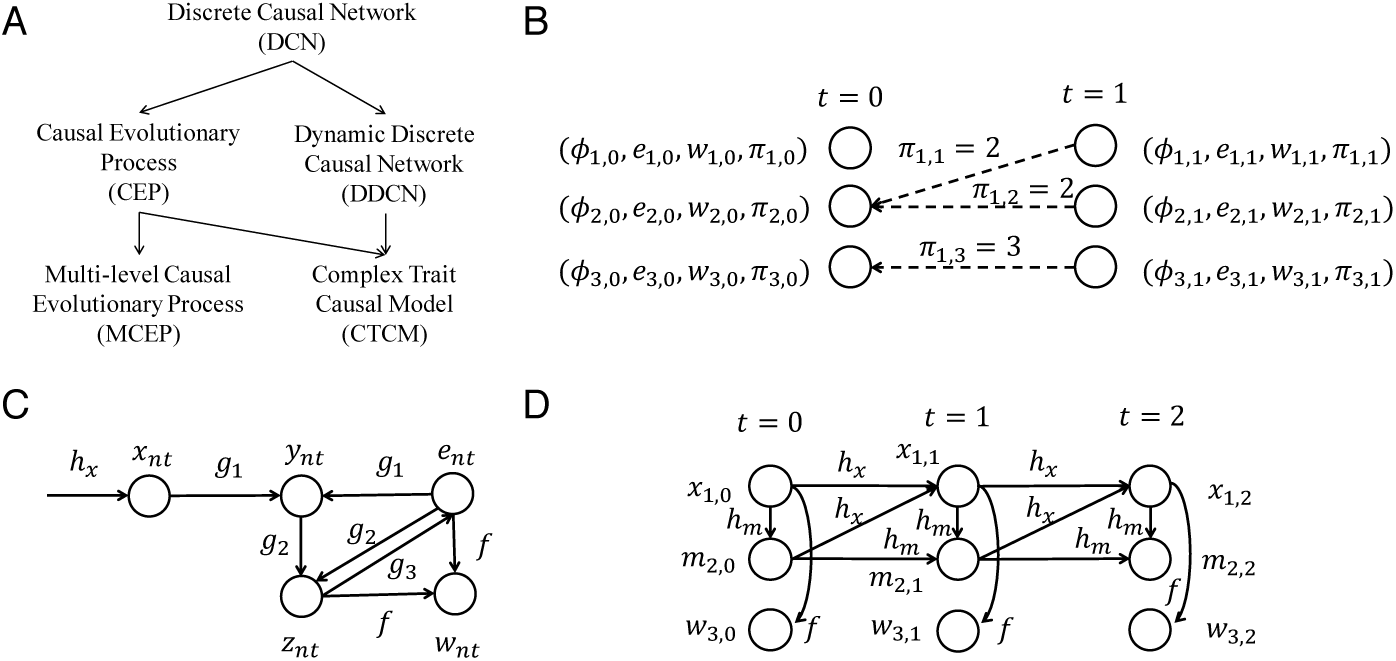
Summary of models. (A) shows the relationships between the main models defined in the paper, where the arrows point from larger to smaller model classes, the latter being a special case (subset) of the former (see Appendix A for DCN and DDCN definitions). (B) shows a schematic of a Causal Evolutionary Process, showing a population of size *N* = 3 at two time steps (nodes representing all variables associated with an individual at time *t*). *ϕ, e, w* and *π* represent phenotype, environment, fitness and parental map respectively, where the latter is also represented explicitly by the dotted arrows, which connect each individual at *t* = 1 to its parent at *t* = 0. (C) and (D) illustrate the CTCM* and MCEP_mut-proc_ models described in Secs. 4 and 5 of the paper respectively. The first is a model of a complex trait, with genetics (*X*), epigenetics (*Y*) and feedback between behavior (*Z*) and environment (*e*), while the second is a model of multilevel selection in cancer with genetics (*X*) and mutational processes (*m*). Solid arrows here represent the directed graphical structure of the underlying discrete causal network (the *P a* relation, which is distinct from the *π* evolutionary variable in (B)). Nodes and edges are labeled with variable and kernel names respectively as defined in these models.

### Multilevel Causality

A further limitation common to straightforward applications of Peal’s *do*-calculus is its seeming reliance on a single level of causal analysis. For instance, in analyzing the causes of an action, it seems appropriate to identify causes at multiple levels, such as nerves firing, muscles contracting, psychological beliefs, desires and past learning. Indeed, in analyzing group selection using causal graphs, a recent approach has distinguished between *causal* and *supervenient* relationships between variables, while maintaining a DAG graphical structure overall [28]. Recent approaches have formalized the idea that a causal system can be described at multiple levels, by introducing coarse-graining mappings between causal structures at different scales [33]. Such approaches capture the relation of *supervenience*, since multiple *finegrained* interventions in one model may be mapped to the same intervention in another provided the causal structure is preserved by the mapping.

The possibility of multiple levels of causation is arguably of central importance in evolution. In particular, we argue that multilevel selection should be seen as a special case of multilevel causality, and introduce a ‘Multilevel Causal Evolutionay Process’ model (MCEP) as a general framework for analyzing such processes. In particular, *fitness* is treated as a causal variable at each level of the MCEP, allowing coarse-grained fitness to supervene on lower-level fitness values (and other evolutionary variables). A particular case we consider is cancer evolution. As has been recently demonstrated [1, 8, 35], tumors not only acquire particular sets of mutations (with positive, negative and neutral effects on growth) over their development, but also acquire prototypical *mutational processes*. These processes are caused by factors such as disruption of the DNA repair machinery or other cellular mechanisms such as DNA methylation, or environmental effects such as carcinogens, which cause particular mutations to become more prevalent depending on local sequence characteristics or chromosomal position. The fact that such processes introduce bias into the way variation is acquired in the tumor means that they can contribute towards the tumor’s evolution. At a fine-grained level, it is the individual mutations themselves which are responsible for fitness variations among cells, and the mutational processes are simply a source of variation. We show however, that by considering a multilevel model, fitness across larger time-scales can be driven by a combination of individual mutations and mutational processes, potentially even primarily by the latter as suggested by recent results [8, 23, 35].

### Causal Transformations

The models we develop in response to the above (cyclic and multilevel causation) both rely on the technical apparatus of a *transformation* between causal systems, as introduced in [33]. In general, this can be thought of as a structure preserving map between causal systems, analogous to a homomorphic map between groups or other algebraic structures. A causal transformation requires that variables and interventions in one causal system are mapped to those in another, while preserving all causal relationships in the first system as seen ‘from the viewpoint’ of the second. Like a group homomorphism, a transformation of causal systems is not necessarily one-to-one or onto, and so the mapping may embed the first causal system in the second, or map many variables in the first onto a single variable in the second. The transformations we consider typically correspond to the latter possibility, and thus can be seen as forms of *coarse-graining*. However, it is important to stress that transformations are not limited to coarse-graining relationships, and are a general mechanism for relating causal systems. We explicitly define the notion of transformation we need for the special case of discrete causal networks in Appendix A, Def. 2.2.

### Information in Causal Systems

An advantage of framing evolutionary models in explicitly causal terms is that it becomes possible to make distinctions between different ways in which evolutionary variables may interact, which are difficult to make otherwise. For instance, it is common to trace variation in a particular trait (such as height) to genetic and environmental sources. With genetics, the (broadly justified) assumption is made that variation in the trait is in response to genetic variation, and thus intervention is not required to assess the causal impact (having controlled for confounders such as population structure). However, the situation is less clear when variation at other levels such as epigentics (transcriptomics, DNA methylation), or environmental factors are considered in relation to high-level trait variation (height, depression). Here, we would like to be able to trace variation to sources which may be involved in cyclic interactions. In general, interventions may be required to assess the causal impact of one variable on another, but it may also be possible to combine observations and assumptions to infer aspects of the causal structure.

For this reason, we also consider how to define a general notion of causal impact in cyclic causal systems, by generalizing the Partial Information Decomposition framework (PID, [40]), which cannot handle cyclic interactions, to a Causal Information Decomposition framework (Appendix A, Def. 2.4). We show that bounds may be derived in this framework that potentially allow direct causal relations between variables to be inferred from observational data by observing non-zero unique information, and differences in observed and causal information between variables to be predicted given assumptions about feedback and interference. We provide technical background for these bounds in Appendix A (Theorems 2.10 and 4.3), draw connections with alternative definitions of causal impact and related bounds (Prop. 2.5), and summarize the implications as they apply to models of epigenetics and multilevel selection in the relevant sections (Th. 4.3 and Prop. 5.5). These results are intended both to motivate the application of models using PID and CID frameworks in analyzing data, and also contextualize the issues underlying existing approaches, even when not coached explicitly in information-theoretic terms.

One application may be in decomposing the genetic, epigenetic and environmental causal factors underlying observed traits, for instance, psychological traits such as depression in the example mentioned above. In this context, methods such as TWAS [41] and related non-linear models [39] aim to isolate the genetic component which is causative for a given trait (for the latter, as mediated by gene expression). However, a more complete picture of underlying causation for a given trait would consider the complete causal effect of (say) gene expression on a trait, and not only the component which mediates genetic risk (since therapeutic interventions are not limited to targeting mediated genetic effects). Environmental variables may also play a role, whose effects may be mediated by epigenetic factors, be independently mediated, or be reflective of rather than causal for a trait (for instance, medication effects). The PID provides a guiding framework for partitioning trait-relevant information between genetics, epigenetics and environment (say). However, it cannot distinguish between causal feedback and feedforward influences of variables on the trait, which is our motivation for introducing the CID; our Theorem 4.3 characterizes the discrepancy between these two frameworks, and thus places a limit on how far the PID can fail to estimate the CID, which (as we show) is characterized by the relative strength of feedforward and feedback processes. While precise estimates of the quantities involved may be difficult, it may be possible to provide plausible empirical approximations; for instance, the feedforward causal epigentic component is bounded by mediated genetic effects, and the feedback component from medication effects can be estimated using animal models, as in recent studies of psychotic medications [4, 11]. In Sec. 5 we also discuss an application in cancer genomics, where we argue that mutation-process specific effects on subclonal fitness may be identified using non-zero mutual information between such processes and growth rate (which can be estimated from sequencing data, as in [34]) via Th. 2.10 and Prop. 5.5.

### Deterministic, Stochastic and Causal Models

Finally, we wish to emphasize the intrinsic differences between the mathematical structures underlying deterministic, stochastic (or probabilistic) and causal models. These kinds of models can be seen as strictly nested inside one another: deterministic models are simply stochactic models whose probabilities are all taken to be either 0 or 1, while stochastic models are causal models which are not subject to any interventions (i.e. subject to the null intervention). In this sense, causal models contain strictly more information than stochastic models, since they represent a *family of distributions* parameterized by all possible interventions, rather than a single distribution. Alternatively, we can say that causal models contain counterfactual as well as probabilistic information. As stressed in axiomatic accounts of evolution [32], we view causal structure as intrinsic to the definition of an evolutionary process, and thus causal models as the appropriate mathematical structure for a complete description of such a process. In Sec. 6, we briefly consider this viewpoint in relation Price’s Equation and a related information-theoretic result based on the Kullback-Leiber divergence (used in information theory as a quasi-distance measure between probability distributions, here the trait distribution at two time-points), discussing their analogues in stochastic and causal models and stressing how the causal view-point offers a more complete picture (albeit, one implicit in other modeling frameworks).

## 3 Causal Evolutionary Processes

We begin by introducing a general model of a *causal evolutionary process*. In its general form, the model is a formalized ‘phenotype-based theory’ of evolution (see [32]), which is agnostic about underlying mechanisms. As argued in [32] (and as we will be elaborated in subsequent sections), such a perspective naturally embeds traditional population genetics models as a special case, since genotypes may be treated as special kinds of discrete phenotypes, while offering a more general viewpoint. For notational convenience, we introduce all definitions and examples below in the context of an asexual population of constant population size, although the model naturally generalizes to mating populations and varying population sizes. Fig. 1B illustrates the model definition below.

### Definition 3.1

(Causal Evolutionary Process (CEP)): *A CEP is a Discrete Causal Network (DCN) over the variables ϕ*_*nt*_, *e*_*nt*_, *w*_*nt*_, *π (all variables discretized, and π*_*nt*_ ∈ {1…*N}), representing the phenotype, environment, fitness and parent of the n’th individual in the population at time t respectively, where ‘fitness’ and ‘parent’ are to be understood in a structural sense to be defined, and n {*1…*N}, t ∈ {*0…*T}*. *We write ϕ*_*t*_, *e*_*t*_, *w*_*t*_, *π*_*t*_ *for the collective settings of these variables at t, and for convenience use identical notation for names and values taken by random variables. Further, ϕ*_*nt*_ *and e*_*nt*_ *may be viewed as a collection of sub-phenotypes and sub-environmental variables, in which case we write ϕ*_*nst*_ *and e*_*nst*_ *for the value of sub-phenotype (resp. environment) s of individual n at time t. We set Pa*(*ϕ*_0_) = *Pa*(*e*_0_) = *Pa*(*π*_0_) = *{} (noting that Pa stands for the ‘graphical parents’ of a variable in the causal graph, while π*_*nt*_ *is the evolutionary variable representing the parent of individual n at time t, whose values are indices of individuals at t* − 1*). For all other variables, we set Pa*(*w*_*t*_) = *{ϕ*_*t*_, *e*_*t*_*}, Pa*(*ϕ*_*t*_) = *{ϕ*_*t*−1_, *e*_*t*_, *π*_*t*_*}, Pa*(*e*_*t*_) = *{e*_*t*−1_, *ϕ*_*t*_, *π*_*t*_*}, Pa*(*π*_*t*_) = *{w*_*t*−_ 1*}*. *A model is specified by defining the following kernel forms, where we use* Π *to denote a ‘kernel product’ (this corresponds to multiplication for directed acyclic causal graphs, but is defined more generally for cyclic graphs as described in Appendix E)*, (=) *for an optional kernel factorization, and set the underlying variables of the DCN to correspond to the lowest-level factorization consistent with the model kernels:*

Fitness kernel:

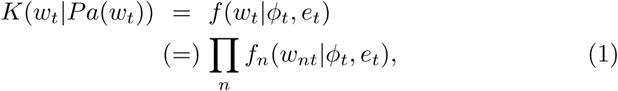

Heritability kernel:

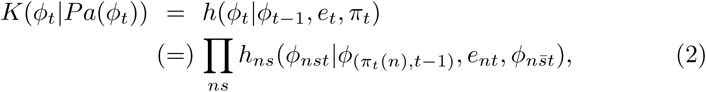

Structure kernel:

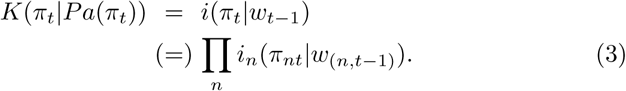

*For the environmental variables, we have the following alternative kernel forms:*

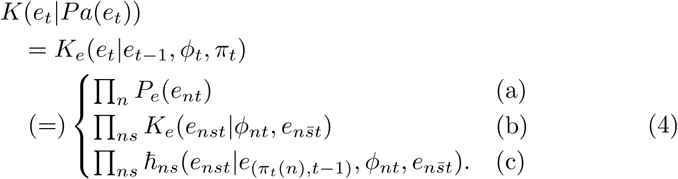

*In Eq. 4, we refer to factorizations (a) and (b) as independent and interactive environmental kernels respectively, and (c) as an environmental heritability kernel (represented by the symbol* n*). Additionally, kernels must be given over the remaining variables for which Pa*(.) =*{} to completely specify a CEP model*.

The interaction of the fitness and structure kernels (*f* and *i* resp.) give rise to different possible fitness models. We summarize some of these possibilities below:

### Definition 3.2

(Fitness models): *We define the following CEP fitness models:* Classical fitness representation:

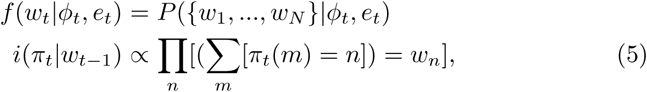

Multinomial model (*ω*_*nt*_ = *w*_*nt*_*/*(∑_*n*_ *w*_*nt*_) denotes normalized fitness):

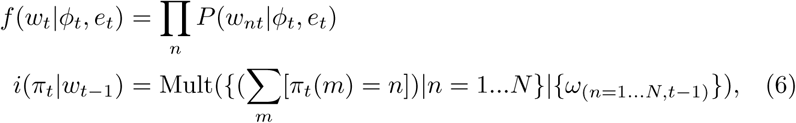

Moran model:

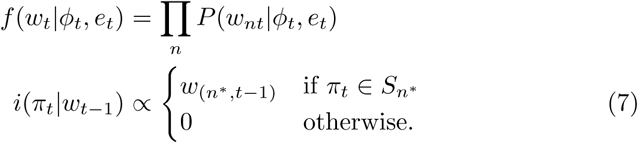

*where S*_*n*\*_ *is the set of all vectors in N*^*N*^ *containing exactly two entries with the value n*^*\**^, *and all other values appear at most once*.

In the classical fitness representation, the variable *w*_*nt*_ directly represents the number of descendants of individual *n* at time *t* in the following generation, and the structure kernel *i* simply ensures the parent map *π* is consistent with these values. In the multinomial model, the values *w*_*nt*_ determine the relative fitnesses of individuals at time *t* and the actual numbers of descendants are determined by multinomial sampling, implemented by *i* (as in the Wright-Fisher model with selection [9]). In contrast, in the Moran model, the *w*_*nt*_’s determine the probability that an individual is chosen to reproduce, and *i* implements the constraint that only one individual reproduces and dies per generation, with the latter being chosen uniformly. Although both Multinomial and Moran models could be represented in the classical fitness representation, this would be at the expense of using a non-factorized form of the *f* kernel; hence we argue that the representations in Eqs. 6 and 7 are more natural parameterizations of these models (so that, in general, ‘fitness’ is not exclusively interpreted as the number of offspring at time *t* + 1, but rather a set of sufficient statistics for generating the parental map at time *t* + 1). Further, we note that, as in the case of the Moran model, the time steps *t* need not correspond to discrete generations.

Finally, we define a *transformation* between CEPs:

### Definition 3.3

(Transformation between CEPs): A *transformation* between CEPs is defined as a transformation between their underlying DCNs in the sense of Def. 2.2 (Appendix A).

In relation to Def. 3.3, we note that a transformation between CEPs need only preserve the *causal* structure; hence, it may (for instance) map environmental onto phenotypic variables, or a large population onto a small population by merging individuals. All that is necessary is that the resulting causal structure may be interpreted as an evolutionary process *in some way*. We shall give examples in the following sections of transformations with characteristics such as above.

## 4 Model 1: Genetics, Epigenetics and Environmental Feedback

The CEP model as introduced in Sec. 3 does not include a model of genetics. However, as noted, the genotype may be regarded as a special type of discrete phenotype, and the process of genetic transmission with mutation can be naturally modeled in the heritability kernel. Here, we describe a type of CEP which includes both genetics and epigenetics, along with potential effects from and impacts on the environment (*environmental feedback*), for instance via behavior or drugs used in treating diseases. The model thus formalizes a *gene-environment-phenotype* map (G-E-P map) of the kind described informally in [16] (for simplicity, we refer to any ‘intermediate phenotype’ as epigenetic, including indicators of cell/tissue state such as the the transcriptome). Our purpose is to provide a general model appropriate for analyzing the causal factors underlying complex traits, such as psychiatric disorders. As we show, using this model, the causal information decomposition (CID) described in Sec. 2 and Appendix A provides a principled framework for breaking down the variation in a complex trait due to genetic, epigenetic and environmental factors; the model is more general than other models with similar goals, for instance TWAS (see [41]), since it aims to model both genetic and other causes of a trait, allowing feedback with the environment at the epigenetic level, and uses an information theoretic framework to decompose the variation and hence is appropriate in the context of arbitrary (non-linear) dependencies.

We first define a general CEP model with the above characteristics:

### Definition 4.1

(Complex Trait Causal Model (CTCM)): *We define a CTCM as a CEP with the following special structure. In terms of phenotypes, we require three sub-phenotypes which we denote X, Y and Z, hence ϕ*_*nt*_ = *{x*_*nt*_, *y*_*nt*_, *z*_*nt*_*}, which represent genotype, epigenome (including transcriptome), and observed trait(s) respectively. Environmental variables are referred to collectively as e. Further, we use the factorized form of the heritability kernel in Eq. 2, and require the following special forms for the sub-kernels (using the kernel product notation from Appendix E):*

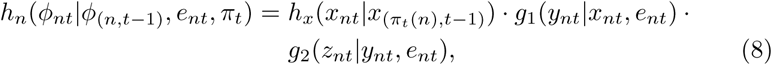

*where g*_1_ *and g*_2_ *are referred to collectively as the* genotype-phenotype map, *and independently as the* genetic-epigenetic *and* epigenetic-observed *kernels respectively, while h*_*x*_ *is referred to as the* genetic transmission *kernel. Further, the CTCM uses an environmental kernel having either an independent factorization, or an interactive form (Eq. 4 (a) and (b) resp.); the latter takes the form:*

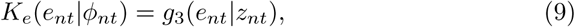

*and g*_1_, *g*_2_ *and g*_3_ *are referred to collectively (when all present) as the* genotype-environment-phenotype map. *[We note that ‘map’ here and following Eq. 8 refers in general to a stochastic map.]*

We next define a special class of CTCMs which embed a DDCM over the *ϕ*_*nt*_ and *e*_*nt*_ variables at each time-point; hence, we model the genetic, epigenetic, observed trait and environmental interactions by an embedded dynamic causal process. We can then coarse-grain this process to a cyclical CTCM over these variables at equilibrium (see Fig. 1C for a related schematic). For conciseness, the full definition of the CTCM* is given in Appendix B, Def. 4.2.

### Definition 4.2

(CTCM with embeded DDCM (CTCM*)): See Appendix B.

The CTCM* model gives us a convenient way of decomposing the variation/information in a trait into components which depend on unique, redundant and synergistic combinations of genetic, epigenetic and environmental factors. Using the CID definition and notation from Appendix A, we propose that, for a population at time *t >* 0, this is achieved by selecting an arbitrary individual *n*, and calculating the backward causal information decomposition *CID*(*S →* <u>*Z*_*nt*_</u>), where *S ⊂{X*_*nt*_, *Y*_*nt*_, *e*_*nt*_*}*, in the eq-CTCM* associated with the original CTCM* (assuming it exists, and that the structure kernel *i* is invariant to permutations of the population indices). Since we specify in Def. 4.2 that the embedded DDCM kernels have the self-separability property, Theorem 2.9 implies that this is a strict decomposition of the variation in *Z*_*nt*_, i.e. *CID*(*S* → <u>*Z*_*nt*_</u>) *≤ H*(*Z*_*nt*_). Further, if *g*_3_ is an independent rather than an interactive environmental kernel, Theorem 2.10 implies that we can lower-bound the forward-CIDs *CID*(<u>*X*_*nt*_</u> *→ Z*_*nt*_), *CID*(<u>*e*_*nt*_</u> *→ Z*_*nt*_), and *CID*(<u>*Y*_*nt*_</u> *→{Z*_*nt*_, *X*_*nt*_, *e*_*nt*_*}*), using the observed unique information between each variable and *Z* (i.e. the observed unique information is predictive of the consequences with respect to an observed trait of performing manipulations on each variable). We note that the PID of the observed distribution in this case is identical to the backward-CID as above.

In the case that *g*_3_ is an interactive kernel, we have environmental feedback from the observed trait to the epigenetic levels, making it harder to relate the observed phenotype distribution to the proposed causal decomposition. However, we can outline a number of possible relationships. For this purpose, we introduce an alternative representation of a CTCM*. We consider that all transition kernels share a common parameter *α* from the self-separable representation, Eq. 28 (with *K*_1_ set to the identity). Since Eq. 28 has the form of a mixture distribution, an equivalent representation of a CTCM* is formed by introducing latent variables, *C*_*Y*_ (*τ*), *C*_*Z*_(*τ*), *C*_*e*_(*τ*), which are Bernoulli variables (or collections of Bernoulli variables if *Y, Z* or *e* are factorized) with mean 1 − *α*. If *C*_*V*_ (*τ*) = 0, variable *V ∈ {Y, Z, e}* does not update at time-step *τ*, otherwise *V* updates according to the *K*_2_ component of the self-separable representation in Eq. 28. If *α* is set large enough with respect to the number of variables, we can ensure that with probability 1 − *ϵ*, with *E* arbitrarily small, ∑_*V*_ *C*_*V*_ (*τ*) *≤* 1, i.e. at most one variable updates at a given time-step. Conceptually, we can view an increase in *α* as effectively a reduction in the duration of the time-step *τ*. We also introduce *C*^*\**^(*τ*) *∈{X, Y, Z, e,∅}*,, writing *C*^*\**^(*τ*) = *V* when *V* is the most recent variable for which *C*_*V*_ (*τ* ^*′*^ *< τ*) = 1 assuming *V* is unique, and *C*^*\**^(*τ*) = *∅* when *V* is not unique. We can then make the following observation (see Appendix B for the proof):

### Theorem 4.3

(Backward-CID bounds): *For a CTCM* represented as above with latent factors C, and associated eq-CTCM*, where S ⊂ {X, Y, e}, V ∈{X, Y, Z, e}*, 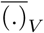 *denotes the mean over values of V, and II is the interaction information (II*(*S*; *Z*; *C*^*\**^) = *I*(*S*; *Z* |*C*^*\**^)− *I*(*S*; *Z*)*), in the limit α →* 1 *we have that:*

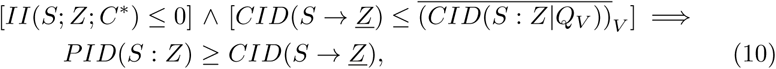

*and similarly:*

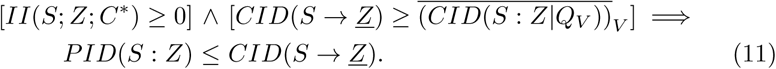

*where all II, CID and PID quantities are evaluated in the eq-CTCM* model (at a given n and t, where C*^*\**^ *is treated as an additional phenotype). Further*, 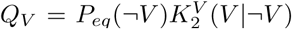 *(unrelated to the notation Q*_*X*_ *used in Def. 2.4) with* 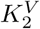 *the second component of V ‘s kernel, as in Eq. 28, and we assume Y, Z and e are not factorized. For the case that Y, Z or e are factorized, S and V are subsets and elements of the sets of relevant factorized variables respectively, and Eqs. 10 and 11 hold identically*.

*Proof* See Appendix B.

Theorem 4.3 shows that we can identify certain situations in which the observed *PID* components at equilibrium consistently under or over-estimate the equivalent components of the *CID*. The LHS of Eqs. 10 and 11 each contain two conditions, the first depending on the sign of an Interaction Information (II) term, and the second comparing two *CID* terms. Broadly, the latter condition implies that if the feed-forward interaction between *S* and *Z* is strong compared to any feedback interactions, the *PID* will tend to underestimate the *CID* (Eq. 11), and vice-versa if the feedback is stronger (Eq. 10). However, the first condition makes this dependent on the type of feedback interactions present: if these tend to interfere with the feedforward interactions so that the net effect is to reduce the mutual information between *S* and *Z* (*I*(*S*; *Z*)), the II term will be positive as in Eq. 11, while non-interfering interactions will tend to increase *I*(*S*; *Z*), potentially leading to a negative II term as in Eq. 10 (note that we use the same sign convention for the interaction information as in [40]). Potentially, the situation in Eq. 11 may apply when *Z* is a disease trait, and *S* is the transcriptome in a relevant tissue, where the feedforward interaction is strong (the trait is strongly determined by *S*), and the feedback (in terms of treatment) reduces the severity of the disease by directly interfering with the underlying mechanisms. In contrast, the situation in Eq. 10 will apply if the feedforward effects are weak, and there is not a strong interference with feedback interactions at the level of *S* (for instance, a disease treatment which targets symptoms in a different tissue, inducing variation in *S* orthogonal to the causal factors for the disease).

## 5 Model 2: Multilevel Selection

Multilevel selection has been identified as an important component in a number of evolutionary contexts, such as eusociality in insects [26], bacterial plasmid evolution [29], and group selection [37]. It has also been proposed that multilevel selection is a driving force behind major evolutionary transitions, such as the transition to multicellularity [6, 27]. Here, we propose a basic definition of a multilevel causal evolutionary process (MCEP) in the framework introduced above, which naturally connects the notion of evolution occurring at multiple levels to coarse-graining transformations in the sense of Defs. 2.2 and 3.3. We then show how two types of multilevel evolutionary process are special cases of our model (group selection, and selection acting on mutational processes in cancer). Our general definition takes the following form:

### Definition 5.1

(Multilevel Causal Evolutionary Process (MCEP)): *An* M-level MCEP *is a collection of causal evolutionary processes, E*_1_, *E*_2_, …, *E*_*M*_, *such that each pair of adjacent processes forms a* 2-level MCEP *in the following sense. CEPs E and F form a 2-level CEP iff there exists a transformation from E to F (denoted* (*τ, ω*)*) in the sense of Def. 3.3, along with a partial map µ of time-points in F to time-points in E (which may depend on* (*ϕ, e, w, π*) *across all variables in E), and the following conditions apply*. **(1)** *We have that* |*F* |_*N*(*t*)_ *≤* |*E*|_*N*(*µ*(*t*))_ *∀t, and* |*F* |_*T*_ *≤* |*E*|_*T*_, *where we write* |*A*|_*N*(*t*)_ *for the ‘actual population size’ of process A at time t, and* |*A*|_*T*_ *for the ‘actual number of time-points’ in process A. Each of these may be different from the values of N and T in A, since we will allow a* null *phenotype value to be declared in each CEP (whose parents are arbitrary, and whose offspring are all* null*): any individuals having ϕ*_*nt*_ = null *will not count towards* |*A*|_*N*(*t*)_, *and time-points for which all individuals are* null *do not count towards* |*A*|_*T*_, *these being the only time-points excluded from the domain of µ; further, time-points beyond* max_*t*_ *{t*|*∃t*^*′*^ *µ*(*t*^*′*^) = *t} do not count towards* |*E*|_*T*_. **(2)** *We require at least one of the inequalities in (1) to be strict*. **(3)** *We require that for any time-point t in E for which µ*(*t*^*′*^) = *t, the projection of the map τ onto ϕ*_*t*_ *in F is not independent of ϕ*_*nt*_ *in E for any (non-null) individual n (i.e. it does not take the same value for all settings of ϕ*_*nt*_ *given a joint setting of all other variables in E), so that no individual’s phenotype is entirely ‘projected out’ at these time-points by τ*.

We now show how the group-selection model of [37] can be represented as an MCEP:

### Example 5.2

(Group Selection model (MCEP_group_)): *We fix an N and T for process E. For all individuals in E, ϕ*_*nt*_ *∈ {C, D*, null*}, where C and D represent cooperators and defectors respectively. e*_*nt*_ ∈ {1…*M} represents the group membership of an individual (where M is the maximum number of groups), and fitness w*_*nt*_ *is determined by the expected pay-off for an individual when interacting with other members of the same group according to a fixed game matrix (see [37]). A maximum group-size is fixed at N*_*G*_, *such that N* = *N*_*G*_*M*. *The heritability and environmental kernels (h and* n*) enforce strict inheritance of phenotype and group membership (with the exception noted below), while structure kernel i implements Moran dynamics (Eq. 7), so long as doing so will not allow a group to exceed N*_*G*_; *otherwise, with probability* (1 − *q*) *a random individual from the same group dies, and with probability q the group divides (implemented by the environmental kernel as a random partition) while all members of another uniformly chosen group die. Since the population number may fluctuate below N*, null *values are used to ‘pad’ the population as required*.

*For process F, we set the population size to be M and the number of time-steps to be T*. *We map t* = 1 *in F to the first time-point in E, and subsequent time-points in F to the times at which the 1st, 2nd, 3rd*… *group divisions occurred in E. The ‘individuals’ in F correspond to the groups in E; hence, we let ϕ*_*nt*_ = (*v*_*n*_, *C*_*n*_), *where v*_*n*_ *is the number of individuals in group n at time µ*(*t*) *in E, and C*_*n*_ *is the proportion of cooperators in group n (we set ϕ*_*nt*_ = null *if v*_*n*_ = 0*). In F, e*_*nt*_ = *{}. We can naturally specify the parental map π*_*nt*_ *on F by mapping a group at t to the group at t* − 1 *when the groups they correspond to at µ*(*t*) *and µ*(*t* − 1) *in E are either the same or split from one another. The fitness model can be specified by letting w*_*nt*_ *be the probability that group n will split first (in the context of all other groups). The structure kernel i then simply needs to implement Moran dynamics, unless the number of groups is less than N*_*G*_, *in which case a* null *group is chosen for replacement in place of uniform sampling. The heritability kernel in F then needs to implement a conditional distribution over ϕ*_*t*_ *corresponding to the joint distribution over the sizes and cooperator prevalences in group at t, given that a particular group from the previous generation divided first (we note that since, in general, dependencies will be induced between the group phenotypes by the intervening dynamics in E, the unfactorized version of the heritability kernel in Eq. 2 must be used). Time-points in F following the last time-point for which µ*(*t*) *is assigned are padded with* null *phenotype values (clearly*, |*F*| _*T*_ *< T, since at most T* − 1 *group divisions can occur in E)*.

*By construction, the pair of CEPs E and F above form a 2-level MCEP*. *For the required transformation, we simply take τ to map a configuration in E to the configuration in F which consistently represents the sizes and proportions of cooperators in each group a at the times µ*(0), *µ*(1)… *by* (*v*_*a*0_, *C*_*a*0_), (*v*_*a*1_, *C*_*a*1_), …. *For the mapping ω we must be careful to restrict the interventions allowed on E to those which fix all phenotypes and environments at a given time t. With this restriction, these can be mapped many-to-one onto interventions in F which match the induced group characteristics. The first two conditions in Def. 5.1 are satisfied by construction, while the third follows since a change in phenotype of a individual in E at a time point µ*(*t*) *necessarily induces a change in the proportion of cooperators in one group, and hence changes ϕ*_*t*_ *in F*.

We note that the MCEP_group_ example above illustrates how the division and complexity of interactions between individuals and environment may depend on the level at which an evolutionary process is viewed (as well as the particular representation): In process *E*, group indices are considered environmental variables, which induce complex inter-dependencies in the fitnesses and heritabilities (phenotypic and environmental) between individuals; however, in process *F*, the groups are themselves considered individuals with their own properties, and much of the complexity at the underlying level is folded into the heritability of group phenotypes, along with a simpler fitness model.

Before outlining our final example, we introduce a general kind of MCEP over multiple temporal levels:

### Definition 5.3

(Regular MCEP with multiple time-scales (MCEP_temp_)): *An MCEP*_*temp*_ *is an MCEP over processes E and F with total time-steps T*_*E*_ *and T*_*F*_, *where T*_*E*_ = *T*_*F*_ *T*_*S*_ *(with T*_*S*_ *>* 1 *a ‘temporal scaling factor’), and µ*(*t*) = *t* · *T*_*S*_. *The transformation τ involves the projection of all phenotype and environmental variables in E onto their values at {µ*(0), *µ*(1), …*}, while the variable π*_*nt*_ *in F is set to the ancestor of n at time-step µ*(*t* − 1) *in E. The fitness variable w*_*nt*_ *in F is set to the absolute number of offspring of n at t* + 1, *and a classical fitness model is used as in Eq. 5. In general, the fitness, heritability and environmental kernels in F will need to take unfactorized forms to capture the complex dependencies induced by the low-level dynamics in E. Further, we restrict interventions in E to interventions on the phenotypes and environments of variables at {µ*(0), *µ*(1), … *}, and map these to corresponding interventions in F*.

We note that any CEP may be converted to an MCEP_temp_ by simply fixing a temporal scaling *T*_*S*_, setting the original CEP as *E*, and following the construction above to form *F*. In this context, the values *w*_*nt*_ represent a limited form of ‘inclusive fitness’ over the period *µ*(*t*) to *µ*(*t* + 1) in *E*, with respect to genealogical relatedness relative to a base population at *µ*(*t*) (see [28] for a discussion of genealogical relatedness and genetic similarity based definitions of inclusive fitness, the former corresponding to Hamilton’s formulation). As our final example, we use the above to illuminate the interaction of mutational processes and selection in cancer. As cancers evolve, subclones acquire not only distinct sets of mutations, but also distinct *mutational processes* governing the random process by which mutations are generated (see [1]). For instance, by disrupting the DNA repair machinery, certain mutations may increase the mutation rate, or make it more likely that specific mutations (e.g. in particular trimer or pentamer contexts) are acquired in the future. Recent evidence has emerged that cancer driver mutations are differentially associated with the presence of particular mutational processes, and that the prevalence of particular mutational processes change in prototypical ways across the development of particular cancers [8, 35]. To analyse the interaction of mutational processes with subclonal fitness, we introduce the following MCEP model (see Fig. 1D for related schematic):

### Example 5.4

(MCEP with mutational processes (MCEP_mut-proc_)): *We build an MCEP*_*mut-proc*_ *model by introducing a CEP model of mutational processes as E, and forming F directly by applying the MCEP*_*temp*_ *definition in Def. 5.3. For E, we set the phenotype variables as ϕ*_*nt*_ = *{x*_*nt*_, *m*_*nt*_ *}, where x and m represents the genotype and mutational processes acting in cell n at time t respectively. We use the factorized form of heritability kernel in Eq. 2, setting* 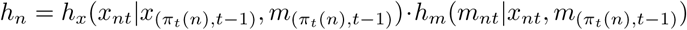. *We note that this incorporates a genetic transmission kernel h*_*x*_ *which is influenced by the mutational processes operating in the parent cell, and a mutational process kernel h*_*m*_ *which allows for potential epigenetic inheritance of mutational processes across generations (as well as determination from the genotype). Further, we use a factorized fitness kernel of the form f*_*n*_(*w*_*nt*_|*x*_*nt*_); *hence we assume that the genotype acts as a ‘common cause’ to the mutation processes and fitness of a given cell, but that the latter two variables are not directly causally linked. The structure kernel can be of arbitrary form, and all environments are empty*.

The MCEP_mut-proc_ example above illustrates the following points. First, we note that for any individual in the lower-level process *E*, the mutational processes *m*_*nt*_ are causally indendent of fitness *w*_*nt*_; that is, intervening on *m*_*nt*_ will not affect *w*_*nt*_ (by definition). However, this is no longer the case in the higher-level process *F* ; here, because of the intervening lower-level dynamics, there is a feedback between the mutational processes and fitness in *E* across multiple time-steps, meaning that *w*_*nt*_ for an individual in *F* is affected by interventions on both *x*_*nt*_ and *m*_*nt*_. In fact, in *F* we have the following:

### Proposition 5.5

(Unique Information bounds for MCEP_mut-proc_): *For an individual n at time t in the high-level component process (F) of an MCEP*_*mut-proc*_ *as above, we have that UI*(*w*_*nt*_ : *x*_*nt*_*\m*_*nt*_) *≤ CID*(<u>*x*_*nt*_</u> *→ w*_*nt*_), *and UI*(*w*_*nt*_ : <u>*m*_*nt*_</u>*\x*_*nt*_) *≤ CID*(*m*_*nt*_ *→ w*_*nt*_) + *C, with C defined as in Th. 2.10. (Appendix A)*

The proof of Prop. 5.5 follows directly from Th. 2.10, along with the MCEP_mut-proc_ definition, which implies that *x*_*nt*_ causally influences, but is not influenced by *m*_*nt*_ (in both *E* and *F*), and both influence and are not influenced by *w*_*nt*_ (in *F*). We note that Prop. 5.5 implies that the unique information components of the observed distribution PID over *x*_*nt*_, *m*_*nt*_, *w*_*nt*_ across multiple generations are informative about the potential contributions of *x*_*nt*_ and *m*_*nt*_ on subclone fitness. Particularly, *UI*(*w*_*nt*_ : *m*_*nt*_ *\x*_*nt*_) *>* 0 implies that there is a generic impact on fitness from the mutational processes across a particular time-scale, providing a lower-bound up to the additive constant *C*. Further, we note that while the MCEP_mut-proc_ model above postulates no intra-generational feedback of the mutational processes on the genotype (only inter-generational feedback), the analysis above can be elaborated to include such feedback within generations, and can be expected to hold so long as the intra-generational feedback is weaker than that across generations.

## 6 Causal Interpretation of Price’s Equation and Related Results

We finish by outlining a number of relationships which can be shown to hold in our CEP framework, including Price’s equation an a number of analogous results. First, we can show:

### Theorem 6.1

(Price’s Equation with probabilistic and causal analogues (assuming perfect transmission)): *In a CEP, with empty environmental contexts and perfect transmission (hence, the heritability kernel factorizes and takes the form* 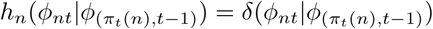, *where δ*(.|*a*) *is a delta distribution centered on a), and a classical fitness model as in Def. 3.2, we have:*

a. Price’s Equation:

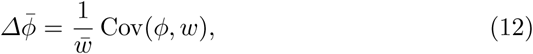
b. Probabilistic Price Equation (Rice’s Equation [6, 32]):

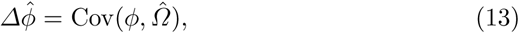
c. Causal Price Equation:

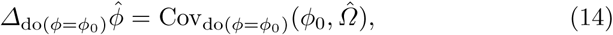

*where we write ā for the average of a across individuals in a single, observed population, and â for the expected average of a across the ensemble of populations modeled by the CEP (following [32])*, 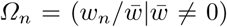 *is relative fitness (see [32]);* Cov(.|.) *is the covariance; and the subscripts* do(*ϕ* = *ϕ*_0_) *indicate that a given quantity is evaluated under the distribution after intervening on ϕ (setting ϕ for all individuals at a given time-step)*.

*Proof* For proofs of (a) and (b) see [27] and [32], which can be applied directly since no interventions are specified. For (c), we note that, having applied the operation do (*ϕ* = *ϕ*_0_), we produce a derived CEP whose underlying distribution is 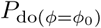. Eq. 14 then follows directly by applying (b) to this derived CEP. □

We note that the distinctions between the original, probabilistic and causal versions of Price’s equation in Theorem 6.1 allow us to make fine distinctions corresponding to direct and indirect selection on traits. For instance, although 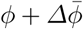 and 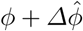 will vary with *ϕ* for any trait which covaries with fitness or expected fitness, for 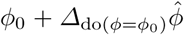 this will only be the case for traits which have a causal impact on fitness (provided the intervention *ϕ*_0_ does not fix all individuals to a single phenotype).

Finally, we note an information-theoretic analogue of the Price equation based on the KL-divergence, which we believe has not been previously observed:

### Theorem 6.2

(Analogue of Price’s Equation based on KL-divergences): *In a CEP with restrictions and notation as in Th. 6.1, we have:*

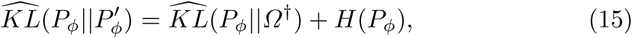

*where P*_*ϕ*_ *and P′*_*ϕ*_ *are the observed (sample-level) distributions across trait ϕ at arbitrary time-points t and t* + 1 *resp*., *KL*(*A B*) =∑ _*i*_ *A*_*i*_ log(*A*_*i*_*/B*_*i*_) *is the KL-divergence between (possibly unnormalized) distributions A and B, H*(.) *is the Shannon entropy, and Ω*^*†*^ *is a vector of relative fitness values for each value of the phenotype*.

*Proof* From the replicator equation, we have:

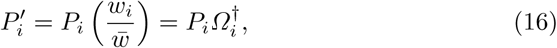

where the subscript *i* ranges across values of the phenotype. The result follows by substituting Eq. 16 into the LHS of Eq. 15 and rearranging:

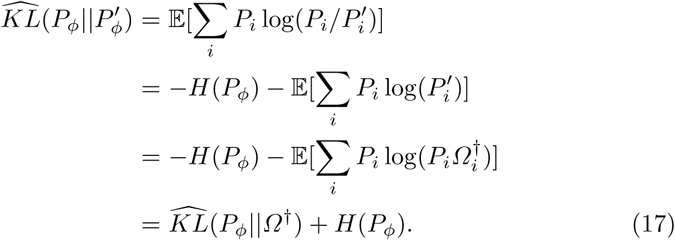

Using a similar argument to Th. 6.1 part (c), we can also state a causal analogue to Eq. 15:

### Corollary 6.3

(Causal analogue of Theorem 6.2): *Using the notation of 6.2, and writing* do(*ϕ*_0_) *for* do(*ϕ* = *ϕ*_0_):

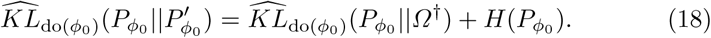

Eqs. 15 and 18 are similar in form to the Price equation, since they determine the distance a trait will move (measured using displacement of its mean or KL divergence over the population-level distribution, for the Price equation and KL-analogue respectively) based on the similarity between the distributions of the trait and relative fitness (measured using the covariance or KL divergence respectively). For complex traits and evolutionary dynamics, the KL-analogues may be more informative, since they model the change in the whole trait distribution, as opposed to only its mean. For instance, at an evolutionary fixed point, we require not only that 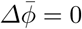 but also 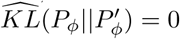 (assuming a large population). We also note the following properties of Eq. 15: Unlike the Price equation, Eq. 15 includes a dependency on the trait’s entropy; further, the KL-’distance’ moved by a trait’s distribution increases as the (unnormalized) KL distance between trait and fitness distributions increases, or the entropy increases; the KL divergence thus need not go to 0 to reach a fixed point, but may be balanced by the entropy term, which may occur since the unnormalized KL divergence is not strictly positive, although it is bounded below by −*H*(*P*_*ϕ*_) since the LHS of Eq. 15 is non-negative (for instance, the uniform distribution on a trait whose values all have equal fitness is a fixed point for which 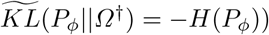. Finally, we briefly note that the relationship in Th. 6.2 differs from the associations between Price’s equation and information theory that have been drawn in [10]: There, the mean change in a trait associated with Price’s equation is re-expressed in terms of the Fisher Information between the trait and the environment (or an equivalent form involving the Shannon information), and it is shown that Fisher’s Fundamental Theorem (FFT) arises by maximizing the information captured by the population; in contrast, Th. 6.2 does not rederive Price’s equation or FFT, but rather relates two KL divergences involving analogous quantities to those in the former, leading to a distinct view of trait evolution at the distribution level as discussed.

## 7 Discussion

The framework of Causal Evolutionary Processes introduced in this paper provides a principled way to formulate evolutionary models, allowing both for cyclical interactions between evolutionary variables, and the analysis of evolutionary processes at multiple levels. We have developed a technical apparatus appropriate for this analysis in the form of Discrete Causal Networks and the Causal Information Decomposition, and have shown how a diverse range of evolutionary phenomena can be captured in our framework, including complex traits produced by feedback processes acting between epigenetic, behavioral and environmental levels, and multilevel selection models, including the selection of mutational processes in cancer. We have explored the properties of these models and our general framework, showing that under certain circumstances the causal impact of a given variable on another (for instance, a variant’s impact on a trait) can be bounded by observed information-theoretic quantities, and that a number of generalizations of Price’s equation hold in our framework.

Our framework may be extended in various ways. For convenience, we have restricted our attention to discrete models in the above analysis (having both discrete time and discrete evolutionary variables). Our current framework may be formulated in the more general context of Cyclical Structural Causal Models [5] (see Appendix C), allowing for a measure-theoretic analysis including continuous variables and time. Further, we have restricted attention to the case of evolutionary processes with asexual reproduction; generalization to processes involving sexual reproduction is straightforwardly handled by altering the structure of the parental map *π* so that individuals are mapped to subsets of individuals in the previous generation as opposed to single individuals, while processes such as recombination and assortative mating can be modeled by using particular forms of heritability and structure kernels.

Further, we have not considered the problem of learning the causal structure and kernel forms from data. Methods relying on Mendelian randomization (e.g. [24]) can estimate the causal effects of variants on a trait assuming a linear relationship, but in general we may be interested in the causal effects of variables above the genetic level (e.g. epigenetics) and environmental factors on a trait, as well as non-linear models. In general, this is a hard problem, but general methods have been proposed, for instance the multi-level approach of [7], or the information-theoretic approach of [19], which may be imported into our framework. Further, we intend the unique information and backward-CID bounds in Th. 2.10 and 4.3 to be relevant for approximating the causal impacts of variables when certain assumptions are made, and in general these may be seen in the context of a host of bounds which relate various kinds of causal effect to observables (without interventions) under a range of assumptions (see [12]).

Finally, we intend our framework to be useful in clarifying foundational conceptual issues regarding causation and evolution. For instance, identifying a realized evolutionary process in nature requires providing criteria for identifying ‘individuals’ on which the process acts, and separating these from an ‘environment’; attempts have been made to cast such criteria in information-theoretic terms (see for instance [22]), and our framework provides a natural language for expressing such ‘connecting principles’. We may for instance declare that, to a first approximation, a single level evolutionary process requires environmental variables which are independent of an individual’s identity given its generation, acting as a ‘thermal-bath’ to the system; features such as spatial population structure, behavior-environmental feedback and niche construction (leading to more complex forms of heritability kernel) would then be taken as second-order principles which, if strong enough, may disrupt the ‘individuality’ of the entities in the original system (and hence the system’s ‘existence’ qua system). Such considerations may also help sharpen questions regarding multilevel selection, whose role has been called into question in explanations of evolutionary processes (see [27, 28] for a summary of the issues). Potentially, a model such as the multilevel CEP we outline, along with principles concerning which types of kernels are more or less ‘preferred’ at each level (for instance, in terms of description length, where lower-level kernels may inherit structure from kernels at higher-levels), could allow us to perform model selection among MCEPs with different numbers of levels. The problem of identifying multilevel selection can thus be cast in the more general framework of identifying causality at multiple levels, where we may have multple levels of variables which supervene on one another (for instance, see [7, 15, 28]); the existence of selection at multiple levels is the particular case of this problem when the causal relationships are constrained to have a special structure (such as an MCEP). In summary, we believe that consideration of explicit causal models of the kind we have outlined will be useful when approaching both computational and conceptual issues in models of evolution.

### Appendix A Discrete Cyclic Causal Systems and Causal Information

We present here a technical summary of the basic concepts we need for our framework. In the process, we introduce the *Causal Information Decomposition* (CID), and summarize a number of its properties. First, we define a model of discrete cyclic causal systems, a *Discrete Causal Network* (DCN), which we will use as our basic model throughout the paper (see Fig. 1A for a summary of the relationships between the main models of the paper). We focus on discrete models to avoid the need to use differential entropy when defining information theoretic quantities, and leave generalization of our framework to the continuous case for future work.

#### Definition 2.1

(Discrete Causal Network (DCN)): *Let 𝒳* = *{X*_*i*_*} be a set of discrete random variables indexed by i ∈ 𝕀* = *{*1…*I}, each taking values in the set 𝒱* = 1…*V, and Pa* : *𝕀 → 𝒫*(*𝕀*) *be a function which returns a set of parents for each index (where 𝒫* (.) *denotes the powerset, and the underlying graph of Pa may contain cycles). Then, a* DCN *over 𝒳 consists of a collection of probability kernels* 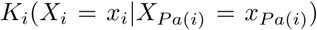 *specifying the conditional distribution of each variable on its parents, and a partial ordering* (*ℐ, ≤*) *where ℐ is a subset of all perfect interventions on 𝒳 with the inherited ordering on interventions (for ι*_1_, *ι*_2_ *∈ ℐ, ≤ ι*_1_ *ι*_2_ *iff ι*_2_ *fixes all variables fixed by ι*_1_ *to matching values, and possibly fixes additional variables). Further, a* solution *to a DCN is a set of joint distributions P*_do(*ι∈ℐ*)_(*𝒳*) *such that the conditional distributions of all non-intervened variables X*_*i*_ *on X*_*P a*(*i*)_ *match K*_*i*_, *and the marginals of all other variables are delta distributions at their respective intervened values*.

We note that our Def. 2.1 can be viewed as a *Causal generalized Bayesian network* as introduced in [17] or a special case of a *Structural Causal Model* (SCM) as in [5], with an additional restriction in each case to a subset of interventions (ℐ, *≤*) (for an equivalent SCM formulation, see Appendix C; also note that for convenience we assume all variables in Def. 2.1 have a common discrete codomain, *𝒱*, which can be assumed without loss of generality, since *V* may be taken large enought to embed all codomains if they are differently sized). By adding the restriction on the interventions considered, we are able to define a notion of *transformation* between DCMs, following the notion of transformations between Structural Equation Models in [33]:

#### Definition 2.2

(Transformations between DCNs): *Suppose we have two DCNs over variables 𝒳* = *{X*_*i∈{*1…*I}*=*𝕀*_*}, 𝒴* = *{Y*_*j∈{*1…*J}*=*𝕁*_*} taking values in {*1…*V*_1_*}*^*I*^ *and {*1…*V*_2_*}*^*J*^ *respectively, with kernels* 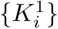*} and* 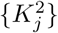 *(resp.) and intervention posets over ℐ and 𝒥 (resp.). Let τ be a map from* 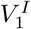 *to* 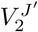, *where J* ^*′*^ = |*A*| *for A* ⊆ 𝒥, *and ω be an order preserving surjective map from ℐ to* 𝒥,. *Then* (*τ, ω*) *is a* transformation of DCMs *iff there exists a pair of solutions for which:*

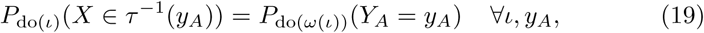

*where τ* ^*−*1^(*y*) *is the pre-image of y under τ*.

Further, we wish to be able to analyse the information shared between DCN variables, and how this is affected by interventions. For this purpose, we first summarize the *Partial Information Decomposition* (PID) framework, using for convenience the formulation in [13]. As originally formulated [40], the PID decomposes the mutual information between a set of predictors *X*_1*…I*_ and a dependent variable *Y*, such that every collection of subsets of predictors is assigned an amount of (non-negative) *redundant* information. As noted by [13], this is equivalent to defining the *union information* for subsets of predictors, which we summarize as:

#### Definition 2.3

(Partial Information Decomposition (PID)): *Given random variables 𝒳* = *{X*_*i*_*} indexed by i ∈* I = *{*1*…I}, with joint distribution P* (*𝒳*), *a collection of (possibly overlapping) subsets {S*_1_, *S*_2_, *…, S*_*J*_ *}, ∀j* : *S*_*j*_ *⊂ I, and a subset T ⊂ I disjoint from all S*_*j*_*’s, we use the symbol PID to denote the* union information, *defined as:*

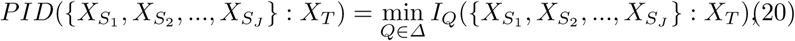

*where I*_*Q*_(*X* : *Y*) *is the mutual information of X and Y under the distribution Q, and Δ is the set of all distributions over {X*_*S*_1, *X*_*S*_2, *…, X*_*S*_*J, X*_*T*_ *} whose pairwise marginals over {X*_*S*_*j, X*_*T*_ *} match those of P, i.e. Q*(*{X*_*S*_*j, X*_*T*_ *}*) = *P* (*{X*_*S*_*j, X*_*T*_ *}*), *∀j*.

Following [3], a number of further quantities may be defined in terms of the PID in the case that *J* = 2. These include the *shared* or *redundant information,* 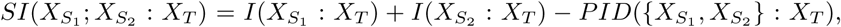, the *co-information* or *synergy*, 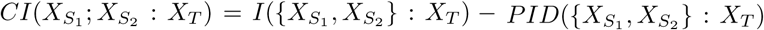, and the *unique information*,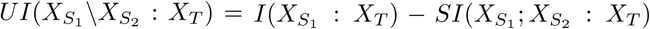. Analogues of these quantities may be defined for *J >* 2 as in [13].

To define a causal analogue to Def. 2.3, we include also a dependency on an *interventional distribution*. Hence, we set:

#### Definition 2.4

(Causal Information Decomposition (CID)): *Given a DCN as in Def. 2.1, subsets over indices {S*_1_, *S*_2_, *…, S*_*J*_ *} and T as in Def. 2.3, and a distribution over interventions, P*_*ℐ*_, *we define the CID as:*

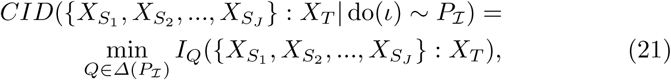

*where:*

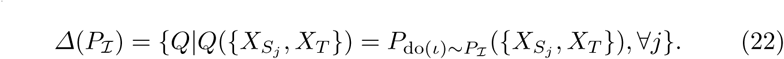

*Further, we use the short-hand notations:*

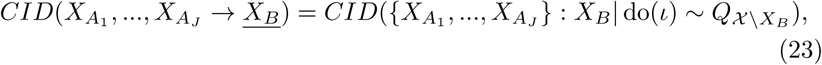

and

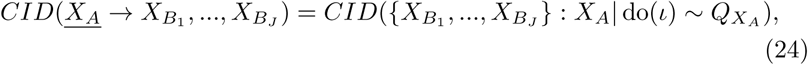

*when ℐ contains a single intervention for each configuration of X*_*B*_ *and X*_*A*_ *in Eqs. 23 and 24 respectively, and* 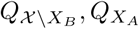 *weight these according to the marginals P* (*X \X*_*B*_), *P* (*X*_*A*_) *(resp.), while assigning* 0 *to all other interventions (hence these intervention distributions capture the* actual variation *in 𝒳 X*_*B*_ *and X*_*A*_ *resp. in the sense of [14]). We refer to Eqs. 23 and 24 as the* backward- *and* forward-*CIDs respectively*.

The CID may be viewed as both a generalization of the PID and the *effective information* (EI) [15, 36]. The EI can be defined as: 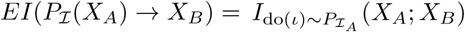, where *X*_*A*_ and *X*_*B*_ are disjoint, and 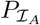 is an interven-tion distribution over *X*_*A*_ (i.e., for any intervention *ι* affecting a variable in 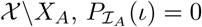). We thus have:

#### Proposition 2.5

(Basic CID identities): *Letting* 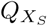 *be as in Def. 2.4, and writing ∅ for the null intervention with δ*_*∅*_(*·*) *the intervention distribution which places probability* 1 *on ∅:*

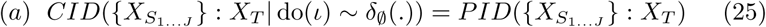

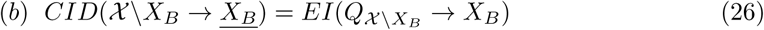

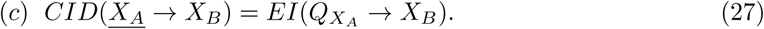

*Proof* For (a), setting *P*_*ℐ*_ = *δ*_*∅*_(*·*) in Eq. 22 makes *Δ*(*P*_*I*_) identical to the set *Δ* in Eq. 20, and hence the identity follows. For (b) and (c), when *J* = 1 in Eqs. 23 and 24, the CID reduces to the mutual information between variable subsets under the intervention distributions 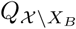 and 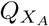 respectively. Hence, setting the EI intervention distributions identically leads to the proposition.□

We now introduce a particular DCM model which will be important in later sections. This is a causal analogue of a Dynamic Bayesian Network [20], which we refer to as a Dynamic DCN (DDCN):

#### Definition 2.6

(Dynamic DCN (DDCN)): *A dynamic DCN is a DCN whose variables and kernel functions have a restricted structure. Particularly, we have 𝒳* = *{X*_(*i,t*)_*} where i ∈ {*1*…I} and t ∈ {*0*…T}, so that X*_(*i,t*)_ *represents an observation of a quantity i at time t. Also, for all t >* 0, *Pa*(*i, t*) = (*Pa*^*′*^(*i*), *t −* 1) *and* 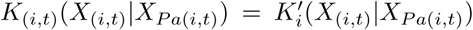, *where Pa*^*′*^(*·)* 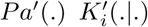 *are auxiliary functions, and for t* = 0, *Pa*(*i, t*) = {}. *Further, it will be useful to consider a restricted DDCN (r-DDCN) with a constrained set of interventions, where we take ℐ to include ∅, along with interventions of the form ι*(*i, v*_*i*_) = do(*X*_(*i,t*=0*…T*_ = *v*_*i*_) *and all combinations of such interventions. For an r-DDCN, we assume that the network converges to a unique steady-state under all interventions. Finally, we define a projected DDCN at time α (p-DDCN(α)) to be a DCN constructed from an r-DDCN, including variables {X*_*i*_*} and {ζ*_*i*_*}, with i ∈ {*1*…I} ranging across the same indices as the underlying r-DDCN, and the X*_*i*_*’s and ζ*_*i*_*’s each taking values from the same set as the X*_(*i,t*)_*’s, augmented in the case of the ζ*_*i*_*’s with a null value* 0. *Writing* (*i*, 0) *for the index of X*_*i*_, *and* (*i*, 1) *for the index of ζ*_*i*_, *we set Pa*(*i*, 0) = (*Pa*^*′*^(*i*)*\i*, 0) *∪ {*(*i* = 1*…I*, 1)*}, Pa*(*i*, 1) = *{}, K*_(*i*,1)_ = *δ*_0_(*′*), *and let K*_(*i*,0)_ *be the conditional distribution of X*_(*i,α*)_ *on X*_(*P a*_ (*i*)*\i,α*) *in the underlying r-DDCN under the intervention ∧*_*i*_*ι*(*i, ζ*_*i*_) *(where ι*(*i*, 0) = *∅, ∀i). The set of interventions in the p-DDCN consists of ∅, along with all combinations of interventions involving* do(*ζ*_*i*_ = *v*_*i*_) *where v*_*i*_ *>* 0. *By construction, the limiting p-DDCN(α) as α → ∞ is well defined, and represents the set of equilibrium distributions (under interventions) of the original r-DDCN, which we denote eq-DDCN*.

We immediately note the following:

#### Proposition 2.7

*For an r-DDCN and a derived p-DDCN(α), we have a transformation of DCNs* (*τ, ω*) *from the former to the latter by setting: ω*(*∧*_*i*_*ι*(*i, v*_*i*_)) = *∧*_*i*_ do(*ζ*_*i*_ = *v*_*i*_), *ω*(*∅*) = *∅ and τ to be the embedding* (*x*_(1*…I,{*0*…T}\α*)_, *x*_(1*…I,α*)_) *↦* (*x*_(1*…I*),*α*_), *where the set A in Def. 2.2 is A* = *{*(*i* = 1*…I*, 0)*}*.

*Proof* The proposition follows directly from the definitions, along with the fact that *ω*(.) as defined is order preserving, since it simply maps the basic intervention *ι*(*i, v*_*i*_) in the r-DDCN to the basic intervention do(*ζ*_*i*_ = *v*_*i*_) in the p-DDCN, implying that order relations between all combinations will be preserved.

Further, we introduce a special separability property on DDCN kernel functions which we will make use of in several places below:

#### Definition 2.8

(Self-separable DDCN kernels): *A DDCN kernel function K*(*X*_(*i,t*)_|*X*_(*P a′* (*i*),*t−*1_)) *will be said to be self-separable, if i ∈ Pa*^*′*^(*i*), *or:*

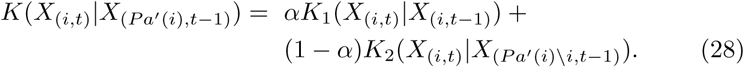

We finish by noting a number of further properties which follow from the definitions above. First, we summarize a number of properties of DDCNs with self-separable kernels as in Def. 2.8:

#### Theorem 2.9

(Properties of Separable DDCNs): *Given an eq-DDCN, derived from an r-DDCN in which all kernels are self-separable, we have (writing H*(.) *for the Shannon entropy):*

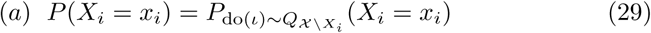

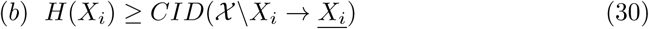

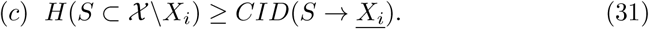

*Proof* For (a), we note that since P(·) is the equilibrium distribution of the underlying r-DDCN, we can write (letting *Y*_*i*_ = *𝒳\X*_*i*_):

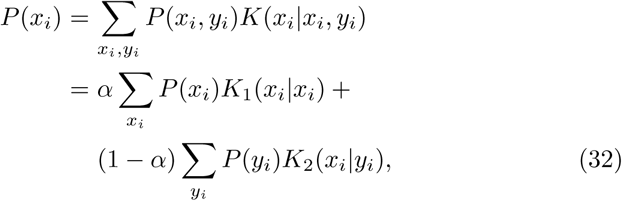

where the second line uses the self-separable property. Since the terms on the RHS depend only on the marginals *P* (*X*_*i*_) and *P* (*Y*_*i*_), the theorem follows, since the marginals over *Y*_*i*_ are preserved in the intervention distribution 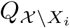

Part (b) follows directly from (a), since *CID*(*𝒳\X*_*i*_ *→ X*_*i*_) is a mutual information involving *X*_*i*_ under the intervention distribution 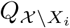. From (a), *H*(*X*_*i*_) is preserved under this intervention distribution, and the mutual information between two variables cannot exceed the entropy of either alone.

Part (c) follows by noting that the entropy *H*(*S*) is also preserved in the intervention distribution 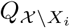. Since the RHS is again a mutual information, the inequality must hold.

We summarize also a number of bounds involving the unique information in DCNs with a more restricted structure, namely a *Pa*(.) function which forms a DAG (i.e. containing no cycles). Particularly, we focus on the effect of an arbitrary variable *X* on another *Z*, where *Z* has no descendants. All other variables are collapsed together as a single variable *Y* = *𝒳\{X, Z}*. Further, we refer to the *causal strength*, ℭ (see [12, 19]), where ℭ_*X→Z*_ = *KL*(*P* (*X*)||*P* (*Y*)*P* (*X*|*Y*)*· P* ^*′*^(*Z*|*Y*)), writing *KL*(.||.) for the KL divergence, and *P* ^*′*^(*Z*|*Y*) = ∑_*x*_ *P* (*X, Y*)*· P* (*Z*|*X, Y*).

#### Theorem 2.10

(Unique information bounds): *For a DCN with Pa*(*·*) *forming a DAG and X, Y, Z as above, we have:*

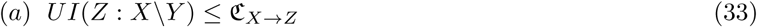

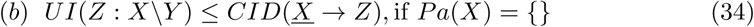

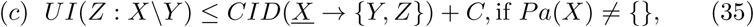

*where*

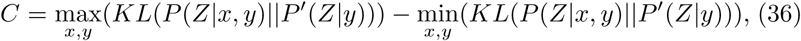

*with P* ^*′*^(*Z*|*Y*) = ∑_*x*_ *P* (*X, Y*)*· P* (*Z*|*X, Y*).

*Proof* For (a), we have from [12] that ℭ_*X→Z*_ *≥ I*(*Z* : *X*|*Y*), where *I*(. :. |.) denotes the conditional mutual information. Further, from [31] we have:

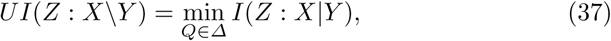

with *Δ* as in Def. 2.3. Since *P ∈ Δ, UI*(*Z* : *X\Y*) *≤ I*(*Z* : *X*|*Y*) *≤* ℭ_*X→Z*_.

For (b), from [12] (prior to Lemma 3), we have that C_*X→Z*_ = *I*(*X* : *Z*) when *Pa*(*X*) = *{}*. Further, for *Pa*(*X*) = *{}* we have that *P*_do(*ι*)*∼QX*_ = *P*, and hence *I*(*X* : *Z*) = *CID*(*X →Z*). Hence, from (a), *UI*(*Z* : *X Y*) *≤*C_*X→Z*_ *≤ CID*(*<u>X</u> →Z*).

For (c), we note that we may write the causal strength as:

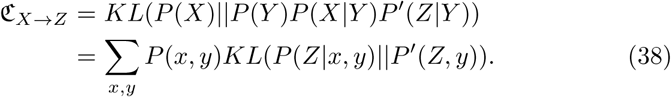

Further, we may write:

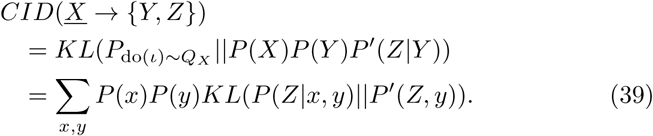

Since Eqs. 38 and 39 are both weighted averages over *∪*_*x,y*_*{KL*(*P* (*Z*|*x, y*)|| *P* ^*l*^(*Z, y*))*}*, we must have:

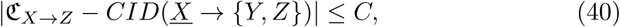

with *C* as in the theorem. The inequality follows from Eq. 40 and (a).

Since the RHS’s of (a), (b) and (c) in Theorem 2.10 may all be regarded as measures of the impact interventions on *X* will have on *Z* (possibly in combination with *Y*), these bounds provide a way of predicting this effect from knowledge of only the observed unique information (i.e. without applying interventions). A corollary of Theorem 2.10 is that if *UI*(*X* : *Z*) *>* 0, C_*X→Z*_ *>* 0, and necessarily *CID*(*<u>X</u> → Z*) *>* 0 if *Pa*(*X*) = {}. We explore Th. 2.10 further through simulations in Appendix D.

### Appendix B Full Proofs and Definitions from Section 4

Below, we give in full the definitions and proofs omitted from Sec. 4.

#### Definition 4.2

(CTCM with embeded DDCM (CTCM*)): *A CTCM* is a CTCM with further structure as follows. We let ϕ*_*nt*_ = *{x*_*ntτ*_, *y*_*ntτ*_, *z*_*ntτ*_ *} and e*_*nt*_ = *{e*_*ntτ*_ *}, where τ is an intra-generational time index, which runs from* 0*…T*_*τ*_. *The kernels of a CTCM* have the form of an embedded DDCM:*

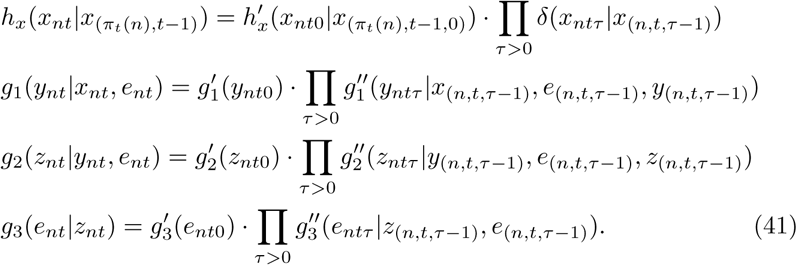

*We allow that these variables and kernels can be further factorized, for instance by decomposing y*_*ntτ*_ *into sub-phenotypes representing expression values of individual genes or gene modules, and e*_*ntτ*_ *into different environmental factors, and introducing sub-kernels of 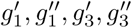 for each sub-variable. Given a lowest level factorization, we require that the all transition kernels (i.e. the kernels 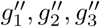, or their sub-kernels) are self-separable in the sense of Def. 2.8, where in all cases K*_1_(.|.) *in Eq. 28 is set to a delta function at the identity (K*_1_(*a a*) = *δ*(*a a*)*). In analogy with Def 2.6, we can define restricted and projected CTCM*’s by applying these constructions to the embedded DDCMs. In the former case, we restrict interventions over the variables X, Y, Z and e to those which fix the variable in an individual at time t across all values of τ, and in the latter case writing p-CTCM*(α) for the CTCM* formed by projecting the phenotype/environmental variables onto τ* = *α at each n and t. By taking the limit τ → ∞, we write eq-CTCM\**=*p-CTCM*(∞). Finally, we note that there is a subtlety in that, in moving from an r-DDCN to a p-DDCN in Def 2.6, we introduce the ‘intervention variables’ ζ; these may be conveniently added as extra environmental variables in a p-CTCM*, since g*_1_, *g*_2_, *g*_3_ *are all conditioned on e*.

#### Theorem 4.3

(Backward-CID bounds): *For a CTCM* represented as above with latent factors C, and associated eq-CTCM*, where S ⊂ {X, Y, e}, V ∈ {X, Y, Z, e}*,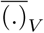 *denotes the mean over values of V, and II is the interaction information (II*(*S*; *Z*; *C*^*∗*^) = *I*(*S*; *Z* | *C*^*∗*^) − *I*(*S*; *Z*)*), in the limit α →* 1 *we have that:*

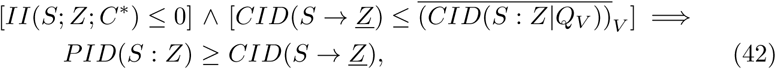

*and similarly:*

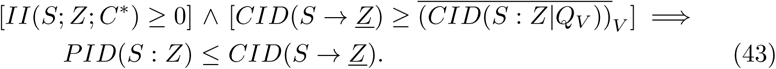

*where all II, CID and PID quantities are evaluated in the eq-CTCM* model (at a given n and t, where C*^*∗*^ *is treated as an additional phenotype). Further*,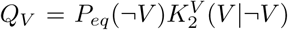 *(unrelated to the notation Q*_*X*_ *used in Def. 2.4) with* 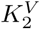 *the second component of V ‘s kernel, as in Eq. 28, and we assume Y, Z and e are not factorized. For the case that Y, Z or e are factorized, S and V are subsets and elements of the sets of relevant factorized variables respectively, and Eqs. 42 and 43 hold identically*.

*Proof* For Eq. 42, we begin by considering the case that, in the underlying CTCM*, at index *τ* we have *C*^*∗*^(*τ*) = *V* ≠ *∅*. Since all other variables *¬V* are arbitrarily sampled and *V* has just updated according to 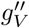 (i.e. letting 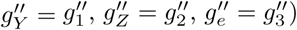), the distribution at time *τ* is *P*_*eq*_(*¬V*)*gV″* (*V* |*¬V*) = *Q*_*V*_. Hence, the mutual information between *Z* and *S* at *τ* is *CID*(*S* : *Z Q*_*V*_). Since we stipulate a common *α* for all transition kernels, the average of this quantity across samples drawn from the equilibrium distribution is approximately the conditional mutual information (neglecting the case in which *C*^*∗*^(*τ*) = *∅*):

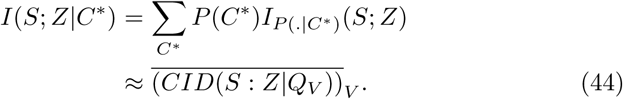

Further, since *II*(*S*; *Z*; *C*^*∗*^) = *I*(*S*; *Z*|*C*^*∗*^) *− I*(*S*; *Z*), in the limit *α →* 1 and for *II*(*S*; *Z*; *C*^*∗*^) *≤* 0 we have:

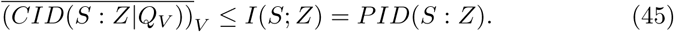

*PID*(*S* : *Z*) *≥ CID*(*S → Z*) then follows from the second line of Eq. 42. For Eq. 43 the proof is similar, with the direction of the inequalities reversed, and the generalization to factorized *Y, Z* or *e* is straightforward.

### Appendix C Representing DCNs as Structural Causal Models

In [5], a *Structural Causal Model* (SCM) is defined as a tuple, *< ℐ, 𝒥, 𝒳, ℰ*, **f**, ℙ_*ℰ*_ *>*, where *ℐ, 𝒥* are finite index sets of endogenous and exogenous variables respectively, *𝒳* =Π_*i∈ℐ*_ *𝒳* _*i*_ and *ℰ* =Π_*j∈J*_ *ℰ*_*j*_ are products of codomains of endogenous and exogenous variables respectively, where each codomain is a measurable space, **f** : *𝒳 ×ℰ →𝒳* is a measurable function, and 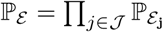 is a product of probability measures over the exogenous variables. A *solution* to an SCM is a pair of random variables (*X, E*) taking values in *X* and *E* resp., such that the distribution of *E* matches P_*E*_, and the structural equations *X* = **f** (*X, E*) are satisfied almost surely.

We may represent a DCN as an SCM as follows (where we assume a DCN solution exists, and construct from this an SCN solution). We let *ℐ* contain indices (0, *i*) for each variable *X*_*i*_ in the original DCN, along with index (1, *i*) for a *mirror variable ζ*_*i*_ corresponding to each original variable (these collectively form the *X*’s of the SCM as defined above). We set the codomains *𝒳*_(0,*i*)_ to be *{*1*…V}* for the *X*’s, and *{*0, 1*…V}* for the *ζ*’s. We then set *𝒥* = *ℐ*_*DCN*_, i.e. the intervention set in the original DCN, and the codomains *ε*_*j*_ are all set to *V* ^*I*^. The probability measure 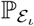 is set so that, if joint configuration [*x*_1_, *…, x*_*I*_] occurs with probability *p* under *ι* in the original DCN, the measure assigned to [*x*_1_, *…, x*_*I*_] under 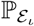 is *p*. We then set **f** so that *f*_(0,*i*)_(*X, ζ, E*) = *v* if *ζ* corresponds to intervention *ι* in the original DCN and *E*_*ι*_(*i*) = *v*; *f*_(1,*i*)_ is the constant 0, and all other values of **f** are set arbitrarily. The intervention *ι* in the original DCN corresponds to making a joint setting of the *ζ* mirror variables in the SCM to the desired intervention values (with 0 corresponding to no intervention). By construction, a joint setting of the endogenous variables surely exists under any intervention in the SCM constructed, since **f** simply ‘copies’ the joint settings from *E*_*j*_ to *X*, where *j* corresponds to the relevant intervention represented by *ζ*.

We note that, in Def 2.1, we use the term *solution* in a slightly different sense to [5]. In our sense, the conditional distributions are specified under each possible intervention, and a set of joint distributions must be found which match these. In [5] however, the full joint distribution over the exogenous variables is specified by the model, and a solution consists of specifying the conditional distribution over the endogenous variables under each possible intervention and setting of *E* which respects the constraints imposed by **f**. Further, we note that while the SCM construction given above is fully general in the sense that any DCN can be represented in the form given, it is also purely ‘formal’ in the sense that the *f*_*i*_’s do not directly correspond to causal mechanisms in the original DCN (represented by the kernels). Clearly, particular DCNs may have more compact representations as SCMs with a stronger correspondence in this sense; for instance, for acyclic DCNs the *ζ*’s are not required, and each *X*_*i*_ may be associated with an *E*_*i*_ *∈* [0 1] which is sampled independently and uniformly, so that *f*_*i*_(*X, E*_*i*_) = *g*_*i*_(*E*_*i*_|*X*_*P a*(*i*)_), where *g*_*i*_(.|.) is the inverse of the cumulative distribution function of the kernel *K*_*i*_(*X*_*i*_|*X*_*P a*(*i*)_), and hence the *f*_*i*_’s correspond directly to the DCN kernels. However, even if such direct correspondences cannot be drawn, the general SCM construction above ensures that for any DCN an SCM exists whose behavior is identical on all interventions.

### Appendix D Simulation study of the Unique Information bound

We explore the behavior of the bound in Th. 2.10 both in conditions when its assumptions are and are not satisfied through simulations. The results are shown in Fig. 2. Here, we run simulations in three DDCN models over the variables *X, Y, Z*, with the connectivity of each model shown on the left (defining the *Pa* map). Each variable can take 4 values (*V* = 4), and we use self-separable DDCN models for all kernels (Def. 2.8). For the kernel parameters, we set *α* = 1 *−* 10^*−γ*^, *K*_1_ to be the identity, *K*_2_ by sampling each transition kernel entry uniformly at random and normalizing so that all conditional distributions sum to one, and we set the initial distributions similarly by uniform sampling. This parameterization lets *γ* act as a ‘stability’ parameter, which we sweep between 0 (low stability) and 5 (high stability), where former implies the identity kernel is never chosen for updates, while the latter implies it almost always is. We first run 10 simulations of each model for *T* = 500 time-steps under no interventions, where a simulation involves sampling the parameters as above, building the full transition matrix *𝖫* over the 4^3^ system states, and analytically calculating *p*_*T*_ = *p*_0_*𝖫T*. From *p*_*T*_, we then calculate all marginal distributions, and use these to calculate *CID*(*V → ¬V*), *CID*(*V*_1_ *→ V*_2_) and *CID*(*¬V → V*) for all variables *V* and variable pairs *V*_1_, *V*_2_ (*V, V*_1_, *V*_2_ *∈ {X, Y, Z}*) in the projected DDCN at time *T* = 500, approximating the equilibrium DDCN (see Def. 2.6). We calculate these quantities by running further simulations under the required intervention models, with the intervention distributions set using the marginals calculated. The latter two quantities allow us to calculate the unique information *UI*(*V*_1_ *\ V*_2_ : *V*_3_) for all variable settings (Def. 2.3 and following). The figure shows, for each variable, a plot which compares the quantity *CID*(*V →¬V*) (the ‘forward’-CID, which may be taken to measure the causal effect of the variable), with the maximum value of *UI*(*V \ V*_1_ : *V*_2_), where *V*_1_, *V*_2_ *∈¬V*. We take the average of these quantities across the 10 simulations for the plots shown.

**Fig. 2.**
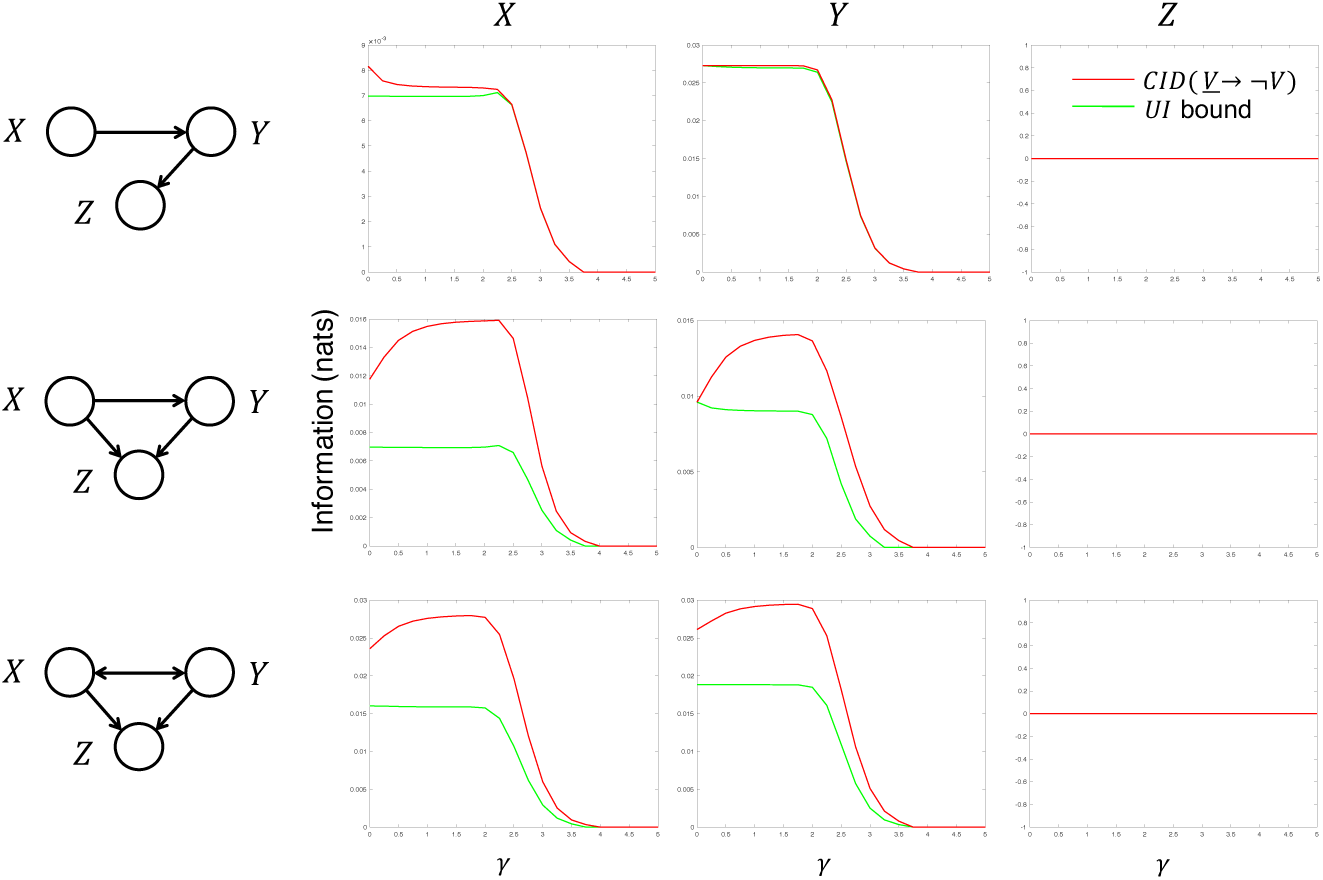
Simulation of the Unique Information bound. Figure shows results of the simulations described in Appendix D. Rows correspond to simulations of the model shown on the left, and columns show the forward-CID (a measure of causal effect) and its unique information lower-bound calculated for each variable. See text for full details.

When the assumptions of Th. 2.10b are satisfied, the quantity max 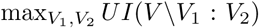 is guaranteed to be less than or equal to *CID*(*V → ¬V*) (and all other unique information bounds will be looser than it). These conditions are only satisfied for variable *X* in the first two models shown. However, variable *Y* in the first two models satisfies the conditions of Th. 2.10c; as shown, the unique information provides a lower bound in these cases also, implying the constant *C* in Th. 2.10c does not typically lead to a violation of the bound under the model sampling distribution described above (e.g. uniform sampling of transition kernels). Further, the last model investigates a case in which the DAG assumption of Th. 2.10 is violated, and we have feedback between *X* and *Y*. The results show that, again, under the model sampling distribution adopted the unique information provides a reliable lower bound here also. We note that in all models, since *Z* has no children, its causal impact (*CID*(*Z → ¬Z*)) is zero; the unique information bound is similarly pushed to 0, hence in the models tested the criterion *UI*(*V \ V*_1_ : *V*_2_) *>* 0 provides a reliable indicator that *V* has non-zero causal impact. The above implies that the unique information bounds of Th. 2.10b and c provide a general indicator of causal impact, which are robust to conditions in which the assumptions of the theorem are not strictly met.

### Appendix E Factorizing kernels in Discrete Causal Networks

We provide here further details on the notation we adopt for factorizations of DCN kernels. As specified in Def. 2.1, a DCN requires a kernel function to be specified for each variable *K*_*i*_(*x*_*i*_|*x*_*P a*(*i*)_) representing the conditional distribution of *x*_*i*_ on its parents. We can summarize a DCN model using a ‘product of kernels’ notation, which we write as either Π_*i*_ *K*_*i*_(*x*_*i*_|*x*_*P a*(*i*)_) or *K*_1_(*x*_1_ |*x*_*P a*(1)_) *K*_2_(*x*_2_ |*x*_*P a*(2)_) *…*. We note that, if the *Pa* relation forms a DAG, this product will directly represent the joint distribution over the DCN variables (subject to no interventions); however, since in general *Pa* may contain cycles, we adopt the convention that Π_*i*_ *K*_*i*_(*x*_*i*_ *x*_*P a*(*i*)_) represents the *set of distributions* which satisfy all the kernel relations. Further, in general, individual kernels may themselves be sets of conditional distributions, although we assume for convenience throughout that the basic kernels use to define a DCN are single distributions (note that a distribution satisfies a kernel only if the conditional derived from the relevant variables matches one in the set associated with the kernel; further, we treat basic kernels notationally as distributions, despite being singleton sets). An intervention which sets *x*_*i*_ to value *v* may be implemented by replacing *K*_*i*_(*x*_*i*_ |*x*_*P a*(*i*)_) by *δ*(*x*_*i*_ |*v*), and a solution to the DCN is a choice function which picks a single distribution from the kernel product sets representing each intervention (including the null intervention). For partial products, this notation represents a higher-order conditional kernel, for instance, consider *K*(*x*_*i*_, *x*_*j*_|*x*_*k*_) = *K*(*x*_*i*_|*x*_*j*_, *x*_*k*_) *· K*(*x*_*j*_|*x*_*i*_, *x*_*k*_). Here, *K*(*x*_*i*_, *x*_*j*_|*x*_*k*_) is a set of conditional distributions over the joint variable (*x*_*i*_, *x*_*j*_), which satisfy the kernel product relations between *x*_*i*_, *x*_*j*_ specified by the lower-order kernels (conditioned on *x*_*k*_). In a given kernel product, we may thus combine groups of kernels together into higher-order kernels, or split them into multiple lower order kernels, while maintaining the solution set for the product. A particular DCN selects a ‘base-level’ factorization, which determines the variable index set I and thus which interventions may be performed on the model; for instance, if *K*(*x*_*i*_, *x*_*j*_|*x*_*k*_) is a base-level kernel, then variables *x*_*i*_ and *x*_*j*_ must be treated as a single variable in the DCN, and interventions cannot be applied to *x*_*i*_ and *x*_*j*_ separately. In this sense, once the base level has been set and a particular DCN solution chosen, all higher-order kernels are fully determined, and are used for notational convenience only.

## Acknowledgment

The authors would like to acknowledge funding from NSF award #1660468. M.B.G. acknowledges support from AL Williams Professorship funds.

